# Novel assay for endangered Hong Kong grouper (*Epinephelus akaara*) to assess eDNA shedding, decay, and population status

**DOI:** 10.1101/2024.03.21.585953

**Authors:** Arthur Chung, Y.C. Kam, Celia Schunter

## Abstract

Overexploitation is a major threat to marine ecosystems, causing collapse of numerous fisheries since the 19^th^ century. The Hong Kong Grouper (*Epinephelus akaara*) is a commercial fish species that suffered at least 50-80% population declines in the past 40 years throughout its distribution range. Yet there has been minimal research or specific management, resulting in insufficient data on abundance, reproduction and habitat utilization. Here we aim to develop a novel species-specific quantitative PCR (qPCR) assay to detect potential occurrence of *E. akaara* through non-invasive environmental water samples. We developed a qPCR assay amplifying 71 bps of the mitochondrial ND2 gene which offers high sensitivity and specificity. To quantify the *E. akaara* population with the emerging environmental DNA (eDNA) tool, however, species-specific shedding and decay rates are crucial. The decay rate of *E. akaara* was similar to that of reported values of other marine fish species. However, the shedding rate of *E. akaara* was found to be few orders of magnitude lower which may be related to the relatively low activity and energy use from solitary and sedentary behavior of groupers. This highlights the importance of empirically determining species or taxon-specific shedding and decay rates to inform accurate abundance estimates with modelling tools for eDNA concentrations. Only 6 out of 88 water samples (6.81%) collected across 4 sampling seasons and 11 sites around Hong Kong showed positive signals at a concentration below limit of detection of the assay, implying its rarity in Hong Kong nowadays. Overall, we demonstrate that eDNA with our qPCR assay is efficient and sensitive in detecting the target species and is a promising tool in documenting endangered species for species management and conservation.

## Introduction

Marine fisheries play a vital role in securing global food supply to provide essential proteins and nutrients in diet (FAO, 2022). Yet, the stability and growth of the industry are facing universal threats by overfishing (Dulvy et al., 2021), in particular from the illegal, unreported, and unregulated (IUU) fishing, which pose significant threats towards establishing sustainable fishery resources (Song et al., 2020). This immense fishing pressure has led to 35.4% percent of fish stocks being fished beyond biologically sustainable levels globally in 2019 (FAO, 2022), and at least 89 globally threatened marine fish species (exclude Chondrichthyes) are still being targeted in industrial fisheries (Roberson et al., 2020). In order to assess the effects of decreasing fisheries resources, it is essential to conduct fish population and stock assessment across multiple spatial and temporal scales. Traditional sampling methods including trawling, gillnets, longlines and traps (Smart et al., 2015), along with fishery-dependent data, such as the volumes of fish catches and landings (Branch et al., 2011) have been frequently utilized to estimate the status of fish stock. Yet these methods generally suffer from various limitations, including high cost, high bycatch rate due to low size and species selectivity, logistic constraints and posed damage to the habitat (Bacheler, 2024). These sampling techniques are also invasive, requiring the capture of fish individuals. This could be extremely difficult for endangered or threatened species due to their low population sizes and could pose further threats to the population.

The emergence of genetic techniques allows for the utilization of environmental DNA (eDNA) as an alternative approach to gather relevant data. This technique is based on organisms leaving traces of their DNA through cellular materials, such as mucus, feces, tissue and blood into the environment that can be harvested and extracted as eDNA (Ficetola et al., 2008; Taberlet et al., 2012). The application of eDNA has benefited fisheries management in terms of rapid and accurate species detection, allowing evaluation of fish species on fishing vessels, live shipment and fish landings to screen for target species e.g. species with high invasive potential and rare/ endangered species (Pikitch, 2018; Roy et al., 2018; Willette et al., 2021). Additionally, it can offer further understanding on the fundamental ecology of commercially important species such as identifying precise spawning grounds and season of catadromous Japanese Eel (*Anguilla japonica*) using qPCR probes (Takeuchi et al., 2019). Such eDNA metabarcoding techniques can reveal fish species community composition with higher species richness and finer spatial heterogeneity as compared to traditional fishing gear sampling techniques (Gehri et al., 2021; He et al., 2022), thus providing more inclusive data to inform fisheries management.

Even though eDNA detection surpasses traditional fishery monitoring methods in terms of strength, it still encounters obstacles when applied to fisheries management. Since eDNA techniques do not involve the processing of fish biological samples, they are unable to furnish specific fishery-dependent data, such as the targeted fish stock’s age, size, weight, life stage, and fecundity (Hansen et al., 2018). Crucially, eDNA does not offer direct insight into the fish stock’s quantity, even though estimating population abundance is a primary goal for fishery management (Lacoursière-Roussel, Côté, et al., 2016). Many factors can influence eDNA detection of a target species, such as organism distribution and density (Eichmiller et al., 2014; Ghosal et al., 2018), diet and life stages (Bracken et al., 2019; Klymus et al., 2015), body size and its relation to metabolic rate (Maruyama et al., 2014a), variation in intraspecific eDNA production associated with stress (Horiuchi et al., 2019; Maruyama et al., 2014a; Nevers et al., 2018), as well as abiotic factors such as salinity, water temperature, solar radiation, pH and microbial activities (Barnes et al., 2014; Lacoursière-Roussel, Rosabal, et al., 2016; Strickler et al., 2015). As such, other than biomass, eDNA concentration is also influenced by complex direct or indirect interactions between multiple biotic and abiotic factors that determine the ultimate shedding and decay rates of the organisms (Hansen et al., 2018; Rourke et al., 2022). Nonetheless, there is a positive relationship between eDNA concentrations and target species abundance or biomass in 90% of 63 relevant studies between 2012 and 2020 (Rourke et al., 2022). Examples include threatened species such as the commercial important Atlantic cod (*Gadus morhua*; Vulnerable) and *A. japonica* (Endangered), where significant correlations were found between eDNA concentration and biomass data from conventional method such as trawl catches and electrofishing (Itakura et al., 2020; Salter et al., 2019).

Despite the high complexity and variability in eDNA dynamics, one way to improve the explanatory power of eDNA is to develop prediction models on eDNA spatio-temporal persistence. Specifically, understanding the roles of environmental conditions in influencing the shedding and decay of eDNA spatially and temporally could predict the fate of DNA materials in different environments and species contexts (Harrison et al., 2019). Previous studies have investigated the shedding and decay of eDNA by mesocosm experiments through introduction and removal of target species to measure persistence time of eDNA, and estimate decay rates through first-order monophasic or multiphasic decay models to demonstrate species and/or system-specific shedding and decay rates with relation to biomass and temperature (Bylemans et al., 2018; Harrison et al., 2019; Jo et al., 2020; Kirtane et al., 2021; Nevers et al., 2018; Sassoubre et al., 2016). These results had demonstrated the importance to empirically determine eDNA shedding and decay rates of the species of interest in different environmental settings which can provide better implications for eDNA data interpretation and build informative eDNA persistence models.

The Hong Kong grouper (*Epinephelus akaara*) used to be one of the most common and dominant grouper fisheries within its limited distribution range from Southern China to Southern Japan in the Northwest Pacific (Figure 1; Randall & Heemstra, 1991). However, its population faced major decline of more than 90% between 1960/70s and 1990s as shown in landing data due to severe overfishing and overexploitation of juvenile stock for capture-based aquaculture (Liu & Sadovy De Mitcheson, 2009). As such, *E. akaara* was listed as Endangered (EN) in the IUCN Red List (Sadovy de Mitcheson et al., 2020), yet still received limited or no species-specific fishery management, such as lack of identification and protection of spawning and nursery grounds in most of the countries within its range (Liu & Sadovy De Mitcheson, 2009). A cost-effective and sensitive method to examine the spatio-temporal patterns of remnants of *E. akaara* populations is crucial to re-evaluate the current population status. Furthermore, information on abundance and habitat association will also allow to shed light on the effectiveness of current marine protected areas. The objective of this study is to (1) develop a probe-based species-specific qPCR assay to accurately detect *E. akaara* eDNA with high sensitivity and specificity; (2) determine the shedding and decay rate of *E. akaara* eDNA to support future eDNA-based models of spatial-temporal persistence, and lastly (3) implement the developed assay in Hong Kong to environmental seawater samples to detect temporal and spatial signals of *E. akaara* in the wild.

**Figure 1.**
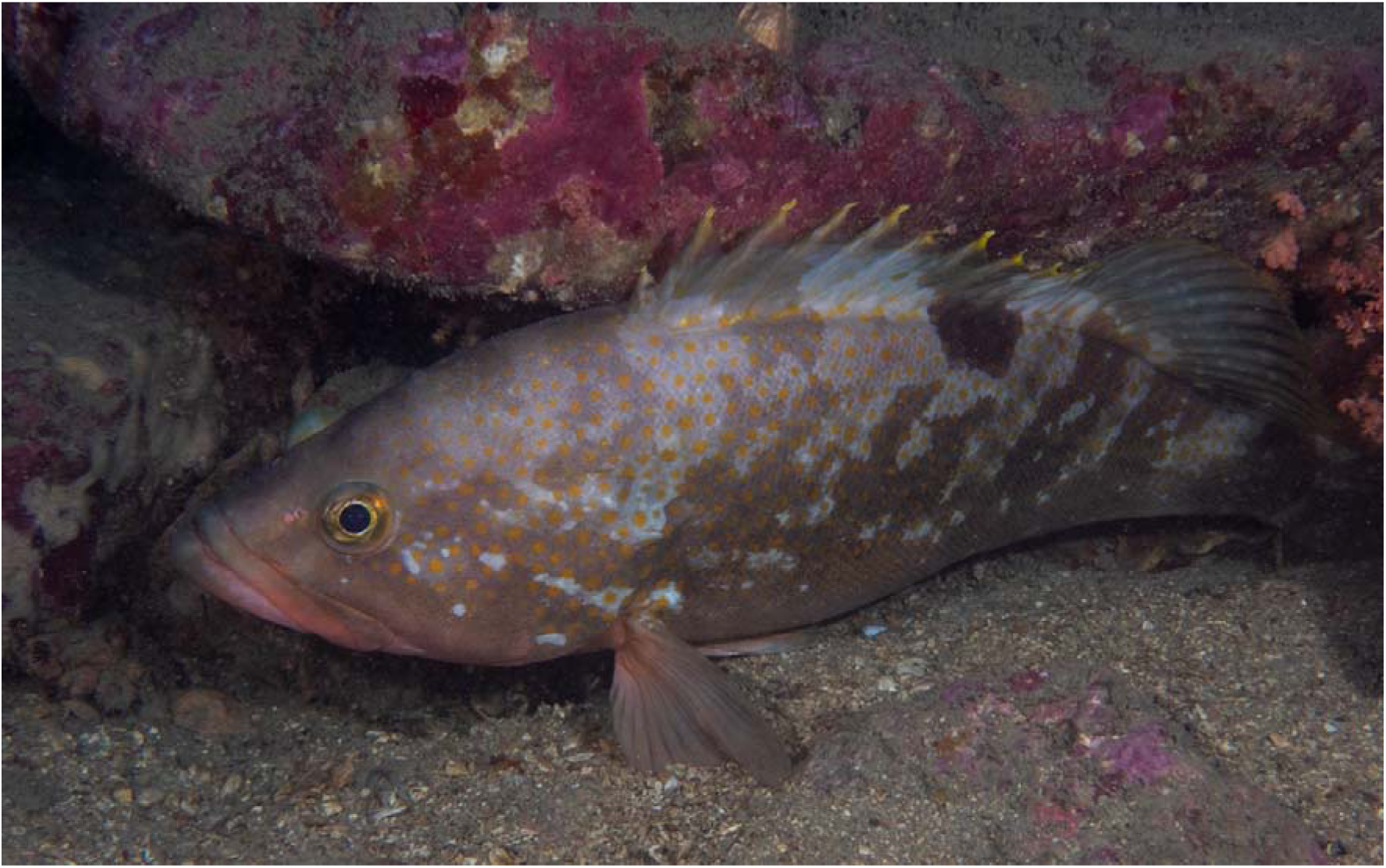
Hong Kong grouper (*Epinephelus akaara*) in its natural habitat. Photograph: YiuWai Hong from 114°E Hong Kong Reef Fish Survey.

## Method

### Development of species-specific assay

To ensure high detection specificity in the assay, we designed a pair of forward and reverse primers that amplify 71 bps of the mitochondrial ND2 gene along with an internal probe that anneals specifically to the target sequence and releases fluorescent signal with the primer design function of Geneious Prime v11.1.4 (Biomatters Ltd.). A reference mitochondrial sequences was first constructed through a consensus alignment from three *E. akaara* mitochondrial available on Genbank (NC_011113, EU043377, KJ700440). To ensure that only *E. akaara* are amplified and not any congeneric species, we obtained mitochondrial sequences on Genbank of species that co-occur with *E. akaara* in the coastal environment of Northern South China Sea (Supplementary figure 1). These were then mapped to the reference consensus sequence to identify and locate gene regions with aggregation of non-matching bases to the reference. On average the number of base pair mismatches to non-target species was 4.39 ± SD 1.58 for the forward primer, 5.06 ± 1.51 for the reverse primer and 3.83 ± 1.20 for the probe. To identify potential secondary structures and avoid dimerization the primer and probe sequences were further analyzed with OligoAnalyzer (Integrated DNA Technologies; IDT). The in silico tested primer sequences (ND2akF: 5’-GCCTTCATTCCTCACAAC-3’; ND2akR: 5’- TAGTGGGAGTAGGAGGATAG-3’) along with the PrimeTime® double-quenched ZEN™/IOWA Black™ FQ labeled with 6-FAM internal probe (ND2akFQ: 5’FAM/TAACACTCCCACTTGCCACTTTAA/3IABkFQ/-3’) were obtained from IDT, Singapore. The specificity of the designed primer sets was tested by running qPCR against genomic DNA (1-3ng) extracted from DNeasy Blood & Tissue Kit from non-target close-related or co-occurring congeneric species that can found in the area (Supplementary table 2). The qPCR reaction mix consisted of 10μl TaqMan™ Gene Expression Master Mix (Applied Biosystems), 500nM of both primers, 250nM probe, 2µl genomic DNA and top up to 20µl total volume with Milli-Q water. The qPCR was run at an initial incubation at 50°C for 2 minutes, then hold at 95°C for 10 minutes, followed by 40 cycles of 95°C for 15 seconds and 60°C for 1 minute respectively. All qPCRs were run with CFX96 Touch Real-Time PCR system (Bio-Rad). Primer/probe sets were considered specific if no amplification (within 40 cycles) at a single-point threshold (100 RFU) was observed for any of the nontarget species genomic DNA.

To determine the efficiency and precision of the developed assay, in particular the limits of detection (LOD) and quantification (LOQ), a curve-fitting method were used. These were based on a 6-point standard dilution curves using a synthetic 300bp DNA gBlock® Gene Fragment as template DNA that covered the entire amplicon region with a starting concentration of 18,000 copies/µl. The definition of LOD and LOQ followed Klymus et al. (2020), where LOD is defined as the lowest concentration at which template DNA produced at least 95% positive replicates, and LOQ as the lowest standard concentration with a coefficient of variation (CV) value below 35%. The modeled LOD and LOQ were calculated using R script provided by Klymus et al. (2020), where sigmoidal modelling is utilized to select the best fitting model for LOD. Among exponential decay, linear, and polynomial models the model with the lowest residual standard error was selected to model LOQ. The figures including calibration curve, plots for both LOD model and LOQ model were also generated from the same script.

### Enclosed aquarium experiment

The experiment to determine shedding and decay rates was conducted in the marine research facility of Ocean Park Hong Kong. To prepare seawater for the experiment, natural seawater was first filtered by sand filters. Water was then pumped to the summit reservoirs with chlorine levels kept at 2-3ppm for disinfection for a minimum of 17 hours. Dechlorination of seawater is done by granular activated carbon filters before transferring the water to the experimental tank.

We minimized the risk of *E. akaara* eDNA contamination before the start of experiment by maintaining the *E. akaara* individuals used in this study in another close loop tank system before the start of the experiment. There was no other *E. akaara* kept in the facility, and no record of wild *E. akaara* individuals nor any fish farms in the source of natural seawater. The 680l tank had a close loop system which circulated for 1 week prior to the start of the experiment, with a chemical UV sterilizer (Tropical Marine Centre) as well as mechanical fibre filters to remove further background eDNA. No target species eDNA was detected through water sampling and qPCR before the start of experiment. The UV filter was turned off throughout the experiment. To determine the shedding rates, two adult *E. akaara*, weighing 0.95kg and 0.90kg, were held in the tank in December 2023 for 72 hours.

The tank was equipped with air bubbling aeration, as well as two powerheads that promoted adequate water circulation within the system. Water parameters including temperature, salinity, dissolved oxygen and pH were measured daily throughout the experiment (Supplementary Table 1). Water collection for eDNA filtration was conducted at timepoint 4.5, 25.5, 29, 47.5, 53.5, 71 hours to calculate the target eDNA shedding rate. To determine the decay rates, water samples were further collected at 7.5, 31, 50.5, and 79 hours after removal of the fish.

For each sample collection point, triplicates of 3L tank water were collected around the periphery at the surface of tank without disturbing the fish that mostly remained sedentary at tank bottom approximate 1m from the surface, and immediately filtered through 47mm polyethersulfone 0.45μm filter (PES; Millipore) with Nalgene reusable filter units (ThermoScientific) using peristaltic pump (Alexis). The filters were kept inside 2ml microcentrifuge tubes, transported in liquid nitrogen and ultimately stored in -80°C until eDNA extraction. New filtered seawater was used to refill the tank to compensate for water loss from sampling and evaporation. A total number of 7 negative controls were filtered on site, which each consisted of 1L of Milli-Q water and were filtered every two days to examine for any contamination.

All laboratory procedures were conducted in molecular laboratory of the University of Hong Kong with standard sterile conditions. eDNA from filters were extracted using the DNEasy Blood & Tissue Kit (Qiagen), with slight modifications from the manufacturer protocol. To the filters 500μl ATL buffer and 30μl proteinase K were added, vortexed for 1 minute and incubated at 56°C overnight. Then, 400μl of the resulting lysate were mixed with 400μl AL buffer and 400μl absolute ethanol with vortex between each addition. The rest of the steps followed the manufacturer instructions until elution, where 100μl elution buffer was added to the spin column and incubated for 3 minutes at room temperature. The quantity and quality of extracted eDNA products were measured by Nanodrop spectrophotometer (ThermoScientific and the DNA was stored in -80°C until further processing.

To determine target eDNA concentration at each time point of the experiment, qPCR reaction mixes and protocols followed the previously described prodecures. No template controls (NTCs), as well as triplicates of standard dilution curve of template gblock DNA were also run together for each qPCR patch to calculate the eDNA concentration of samples in the same batch.

### eDNA decay rate modeling

The experimental tank was modelled as completely mixed batch reactors to determine shedding and decaying rates of *E. akaara* eDNA following previous studies (Kirtane et al., 2021; Sassoubre et al., 2016) with the following equation (Equation 1):

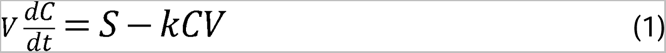

Where *V* is the volume of the tank, *C* is the concentration of *E. akaara* eDNA in copies/L, *t* is time in hours, *S* is the shedding rate and *k* is the decay rate constant. We assume the tank content is well mixed, and the degradation process followed first-order decay with a monophasic decay model (Harrison et al., 2019). The decay rate constant *k* was calculated during the decay period after fish removal by fitting linear regression to ln(*C/C_0_*) versus time (h), where *C_0_*is the steady state DNA concentration. During the eDNA concentration steady state which was achieved after xx hours of the experiment, the change in eDNA concentration is zero (*dC/dt = 0*), thus shedding rate is *S = kCV*. Shedding rate was then calculated from the decay rate constant in copies per hour and kg of fish to adjust for biomass and associated with standard deviations determined by propagating errors for each multiplied term *k, C* and *V* following Sassoubre et al. (2016). Data from the last sampling timepoint, which was 79 hours after fish removal, were removed from decay rate constant calculation as it does not follow the preassumed first-order decay which could due to resuspension of eDNA particles (Harrison et al., 2019). eDNA half-life (T_1/2_) was also calculated following Kirtane et al. (2021) with Equation 2:

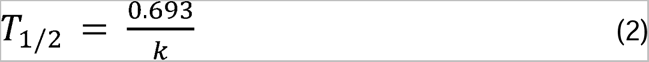

Where *k* is the decay rate constant used in Equation 1.

### Study area and eDNA collection

To implement the developed assay in marine water, we conducted water sampling across Hong Kong temporally and spatially. The sampling was conducted in four occasions, including May, August, November of 2022 and February of 2023 respectively. *E. akaara* in Hong Kong were mainly distributed in the Eastern waters according to previous survey and fishermen records, inhabiting coastal shallow coral and boulder reefs (Liu & Sadovy De Mitcheson, 2009; Sadovy & Cornish, 2000). As such, 11 sites including previous recorded locations and local marine protected areas where both recreational and commercial fishing are prohibited were selected for eDNA collection (Supplementary Figure 2). Two replicates of 30L seawater were collected and filtered on site along a 2km transect along the shore. Water was collected with peristaltic pump (Alexis) on boat at a speed of approximately 5- 6mins/L and water depth of 1m. For each replicate, seawater was first passed through a 10 μM string wound prefilter (Thomson) to remove excessive large particles to minimize filter clogging, which was shown to not affect the number or identity of fish species while reducing PCR inhibitors such as humic acid in the samples (Takasaki et al., 2021), and in turn filtered through a 0.45μm 600cm^2^ PES filter (Waterra). Sterilized tubing and gloves with new prefilter were replaced before the start of filtration in each site. After filtering, each filter was filled with 80ml CTAB lysis buffer and stored in room temperature. DNA extraction were performed on all filters within two weeks of sampling.

### eDNA extraction

The extractions of a total of 88 filters from 11 sites in four different seasons were based on modification of the low-salt CTAB protocol from Arseneau et al. (2017). Each filter was first shaken manually for 1 minute, then the buffer inside was poured out and separated into four replicates of 20ml in 50ml falcon tubes. Two replicates were returned to storage at 4°C, while the other two proceeded with addition of 100µl proteinase K solution (ThermoScientific) and incubation at 60°C for an hour inside a thermal incubator (Eppendorf). After cooling down to room temperature, equal volumes of Chloroform: Isoamyl alcohol (24:1) were added to each tube, mixed by inverting the tubes, and then centrifuged at 3000g. Subsequently, the aqueous phase from the centrifuge product was then mixed with twice the volume of diluted CTAB buffer (CTAB buffer without sodium chloride solution) and incubated at 60°C with 180 rpm for at least one hour until white suspended crystals were observed. The precipitate was collected by centrifuge at 15,000g for 5 minutes, dislodged by 80% ethanol and transferred to a new 1.5ml microcentrifuge tube. The pellet was rinsed twice with 80% ethanol to remove excess CTAB. The tubes were then dried under a fume hood for 10 minutes to remove excess ethanol. Finally, the DNA pellets were resuspended with 60°C 100µl low EDTA TE buffer (Invitrogen), and incubated at 58°C for 15 minutes to dissolve the DNA. The quantity and quality of extracted DNA products were measured by Nanodrop spectrophotometer (ThermoScientific). Replicates from the same filter were pooled and stored in -80°C. qPCR reaction mixes with 5 replicates per filter, protocols, NTCs and standard dilution curves were prepared and run as previously described.

## Results

### Designed assay performance

The designed species-specific assay for *E. akaara* was optimized and had a amplification efficiency of 95.7%. The LOD and LOQ of this assay were determined as 3.71 and 30 copies per reaction respectively, as revealed from the curve-fitting modeling method (Table 1; Supplementary Figure 3; Supplementary Figure 4). No non-specific amplification was found within 40 cycles of qPCR in congeneric species genomic DNA samples.

**Table 1.**
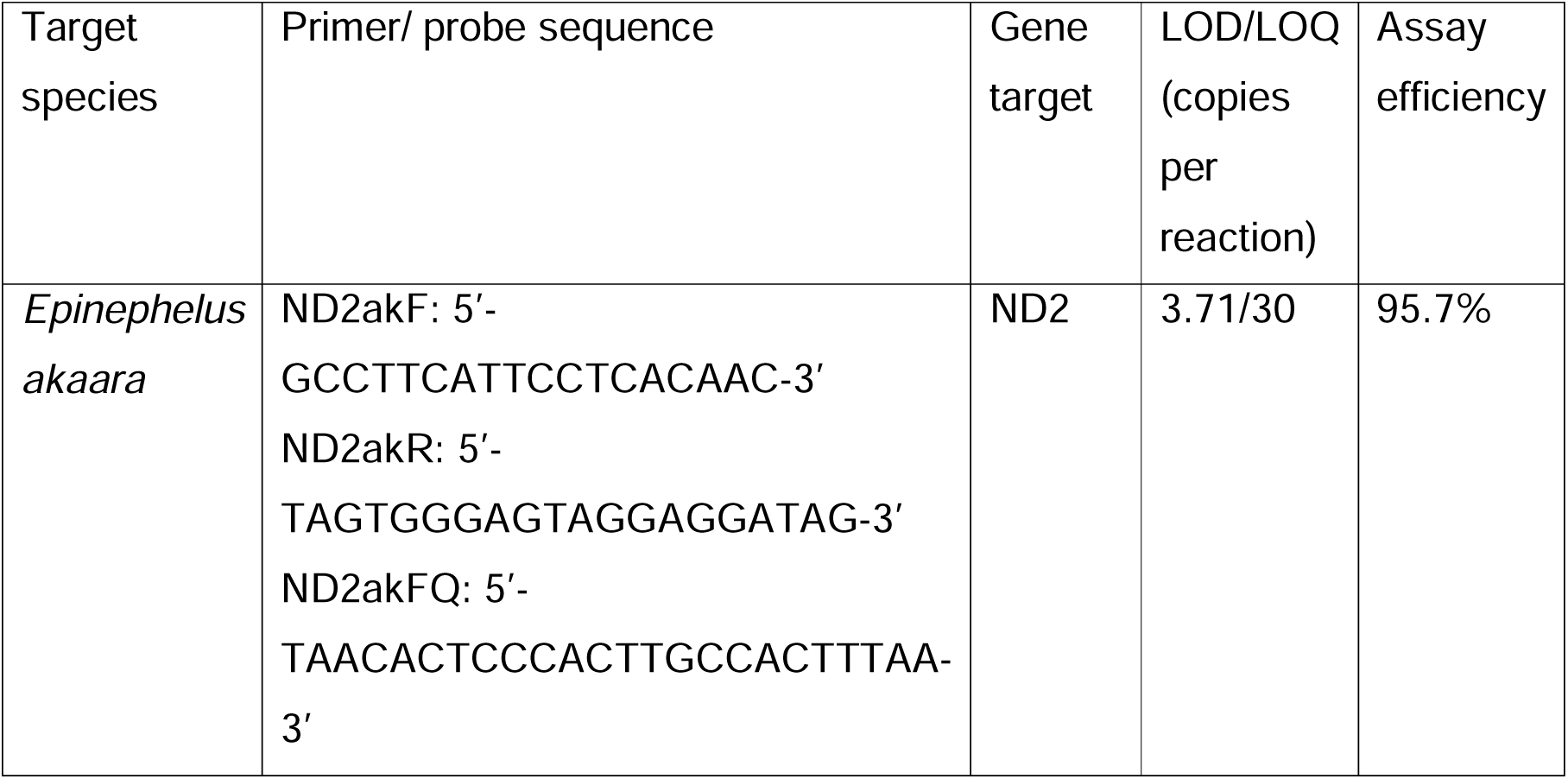
Details of the optimized species-specific primers and probe for *E. akaara* designed in this study.

### Shedding and decay rates of *E. akaara* eDNA

*E. akaara* eDNA concentration quickly accumulated once the two *E. akaara* individuals were introduced into the experimental tank, as seen from the high eDNA copy number per liter at t=4.5 hours (Figure 2). The eDNA concentration remained stable until removal of fish, therefore a steady state of eDNA concentration was assumed and calculated from time period of 25.5 to 71 hours, which was 8284 ± 536 copies per liter. Once the fish were removed, the eDNA concentration dropped immediately and fell below LOQ levels in 50.5 hours. With a constant decay rate of 0.131 ± 0.0111 per hour, the shedding rate was calculated as 3.69 x 10^5^ ± 3.93 x 10^4^ copies per hour individual fish, or as 3.89 x 10^5^ ± 4.14 x 10^4^ copies per hour and biomass in kg (Table 2). The eDNA half life (T_1/2_) was found to be 5.29.

**Figure 2.**
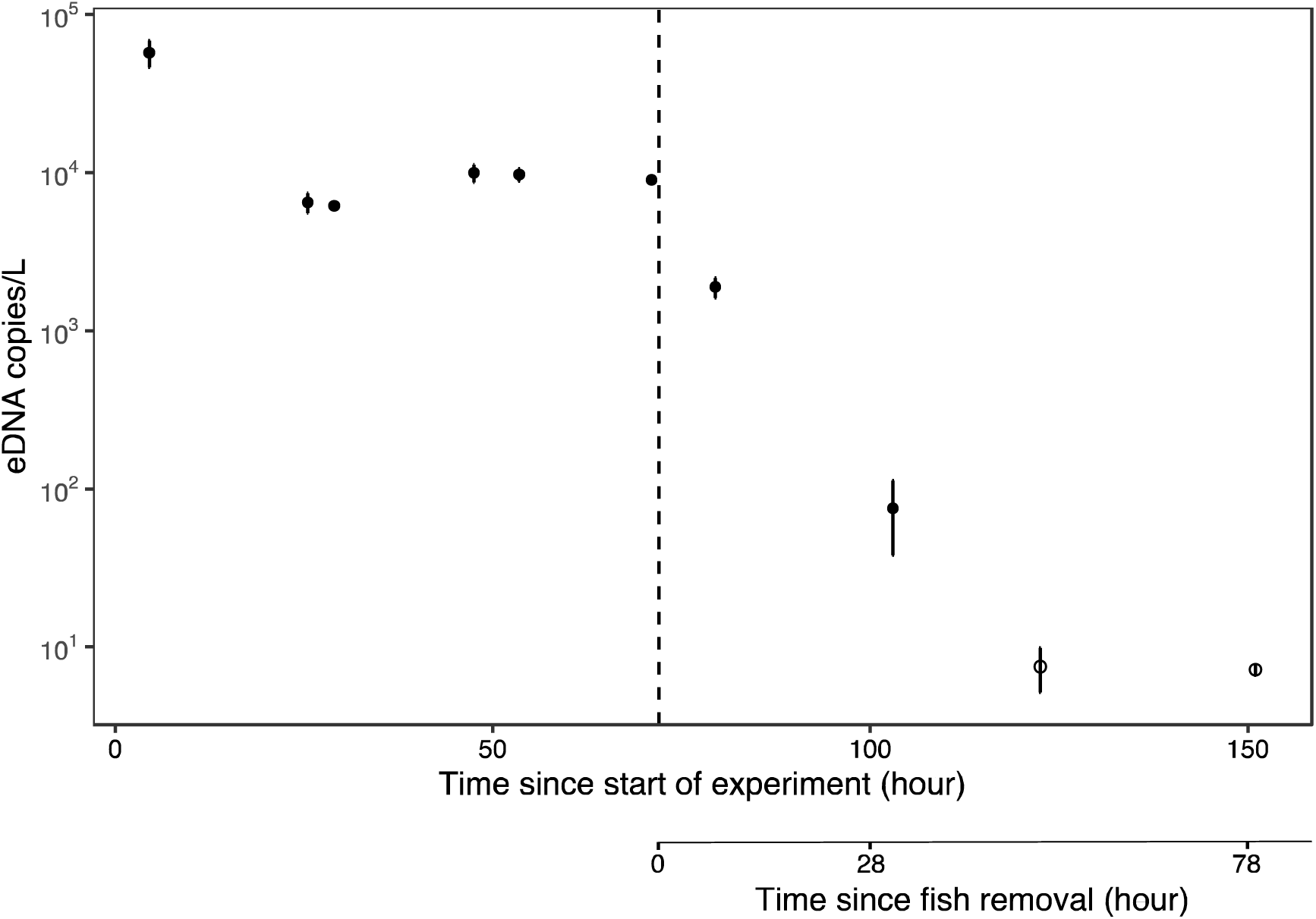
eDNA concentration in copy numbers throughout the experiment. Fish were introduced into the experimental tank at the beginning (time = 0). Error bars represented standard error at each sampling timepoint. Vertical dashed line indicated time of removing the fish from experimental tank. Sampling timepoint with points of open circle indicate eDNA concentrations were found below LOQ.

**Table 2.** eDNA Steady state concentration, shedding rate and decay rate constant from the *E. akaara* experiment.

| | Steady state concentration (copies/ L) * | Shedding rate per fish (copies/h /individual) ** | Shedding rate per kg (copies/h /g) ** | Decay rate constant per hour * | eDNA half life $T_{1/2}$ (h) |
| --- | --- | --- | --- | --- | --- |
| *Epinephelus akaara* | 8284 ± 536 | $3.69 \times 10^5 \pm 3.93 \times 10^4$ | $3.89 \times 10^2 \pm 4.14 \times 10$ | $0.131 \pm 0.0111$ | 5.29 |

\* Errors represented standard error
\*\* Errors represented propagated error

### eDNA detection in environmental seawater samples

There was no *E. akaara* eDNA detected for most collected environmental filtered seawater samples across spatial and temporal scale. For only 6 out of 88 samples (6.82%) minimal eDNA was detected, yet the concentration was very low at a level below the LOD (Table 3). The detection probability from these samples were also very low, as in most cases only 1 out of 5 qPCR technical replicates resulted in amplification. In total, minimal traces of *E. akaara* eDNA were only found in 4 out of 11 different sites across all different sampling seasons except May 2022 (Table 3).

**Table 3.** Details on environmental seawater sample with positive *E. akaara* eDNA detection. Locations of the sampling site are provided in Supplementary Figure 2.

| Date | Sampling site | Number of amplified qPCR technical replicates (out of 5) | Concentration |
| --- | --- | --- | --- |
| 20220818 | Kung Chau (KC) | 1 | Below LOD |
| 20220819 | Che Lei Pai (CLP) | 2 | Below LOD |
| 20221116 | Kung Chau (KC) | 1 | Below LOD |
| 20230228 | Cape d'Aguilar (SWIMS) (filter replicate 1) | 1 | Below LOD |
| 20230228 | Cape d'Aguilar (SWIMS) (filter replicate 2) | 1 | Below LOD |
| 20230228 | Ninepin Island (NP) | 1 | Below LOD |

## Discussion

Development of eDNA tools coupled with empirically derived eDNA production and degradation rate can provide an effective and non-invasive method to detect and quantify rare species. In this study, we successfully developed a highly specific and sensitive qPCR assay for the endangered *Epinephelus akaara*. To our best knowledge, this is the first study examining shedding and decay rates on a species from the family Serranidae, offering valuable data for biomass estimation and fishery stock management for these commercially targeted taxa. No *E. akaara* target sequences were detected in most of the spatially and temporally collected water samples, suggesting the likely absence of this species in the surveyed areas. We demonstrated the high sensitivity of eDNA coupled with qPCR in inferring low abundance or absence of target species.

### *E. akaara* species-specific eDNA assay development

High amplification efficiency, sensitivity and specificity are found in the developed eDNA assay to detect *E. akaara* with custom built primer pairs and internal probes. Specifically, the assay showed an amplification efficiency of 95.7%, which falls within the acceptable criteria range (90%-110%) of qPCR validation guideline set by Broeders et al. (2014), and indicates that a high fraction of target DNAs are copied in each cycle. The curve-fitted model determined limit of detection (LOD) of this assay to be at 3.71 copies per reaction, which is at similar levels to other established eDNA assays on fish species, ranging between 2.19 to 22.1 copies per reaction (Kirtane et al., 2021; Klymus et al., 2020; Pont et al., 2023). On the other hand, the determined LOQ was 30 copies per reaction, which is an order of magnitude lower than some of the established fish eDNA assays at a range of 10 to 1000 copies per reaction (Kirtane et al., 2021), demonstrating the ability of the developed assay to detect target eDNA even at low concentration levels. In addition, the designed assay with maximized mismatches in primer pairs and probes did not cross-amplify phylogenetically closely-related species including *E. awoara* and *E. fasciatomaculosus* (Ma et al., 2016), as well as co-occurring congeneric species in the northern South China Sea, demonstrating its high specificity. Distinguishing closely related species with qPCR assay can present a challenge due to their high sequence similarity which can lead to erroneous signals, either false positives or false negatives (Wilcox et al., 2013). With these successful validations, our developed assay has achieved a level 4 out of 5 according to the validation scale set by Thalinger et al. (2021), which offers specification to classify and evaluate assays on their accuracy and sensitivity for single species detection and their readiness for routine species monitoring. eDNA assays were only ever able to reach level 4 (Thalinger et al., 2021). Overall, we have shown that our developed assay is of high standard providing very low false positive detection probability through rigorous validation.

### *E. akaara* eDNA shedding rate and decay rate

The highest peak of *E. akaara* eDNA concentration was observed at the fist sampling point of the experiment at 4.5 hours, which likely originates from additional shedding of eDNA due to stress from handling as has been seen in previous studies (Kirtane et al., 2021; Sassoubre et al., 2016). The effect of additional input of eDNA was eliminated by allowing for the prolonged acclimation of fish and accumulation of eDNA in the tank, as seen from the slight drop of concentration that quickly reached a steady state until the removal of the fish, allowing the calculation of shedding and decay rate.

The shedding rate of *E. akaara* eDNA in this study was lower than those of other marine fishes previously reported. This is unexpected, as many species showed relatively similar levels of shedding rate (Kirtane et al., 2021; Plough et al., 2021) with a shedding rate per hour per biomass a few order of magnitudes higher than that of *E. akaara* (3.89 x 10^2^ copies/h/g), such as the Black Seabass (5.72 x 10^4^ - 2.79 x 10^5^ copies/h/g) (Kirtane et al., 2021), Pacific Sardine (1275 pg/h/g) (Sassoubre et al., 2016), Atlantic Sturgeon (2067-1331 pg/h/g) (Plough et al., 2021), and the Japanese Jack Mackerel (10^5^ – 10^7^ copies/h/g) (Jo et al., 2019). *E. akaara*, or *Epinephelus* spp. in general are solitary and sedentary species that spend most time in the benthic environment (Gibran, 2007) with limited movement and energy use compared to other species with more mobility or even schooling behaviors. From our experimental observations, *E. akaara* were found to be sedentary on the bottom of the tank and inactive most of the time, which may lead to low fish activity and energy use, and subsequently fewer eDNA being shed (Thalinger et al.,2021) hence relating fish physiology and behavior to eDNA concentrations. We also did not observe any noticeable signs of shed scales nor mucus from *E. akaara* in the tank, which were recorded previously in species with higher shedding rate (Sassoubre et al., 2016), suggesting low rate of body material shedding from *E. akaara* may also account for the low shedding rate. Considering *E. akaara*’s stationary behavior and the fasting throughout the experiment, we believe the reported shedding rate may represent a lower bound for adult *E. akaara*. Overall, our results represent a species-specific eDNA shedding rate for adult *E. akaara* and demonstrate the necessity of determining shedding rate of species of interest, as shedding rate is shown to be species-specific and may vary significantly between species. This also suggests that eDNA concentration will depend on a number of physiological and behavioral factors other than biomass alone.

The first-order decay rate constant for *E. akaara* eDNA (0.131) however is similar to reported values of other marine fish species in previous studies, such as black seabass, winter flounder and summer flounder (0.07-0.57; Kirtane et al., 2021), or slightly higher than species such as northern anchovy, pacific sardine and pacific chub mackerel (0.055-0.101; Sassoubre et al., 2016). The consistent decay rate among various species is likely attributable to the processes of cellular and spontaneous degradation, influenced by factors like the molecular state of eDNA and environmental conditions such as temperature, pH, and solar radiation exposure, rather than solely by the species involved. (Hansen et al., 2018; Harrison et al., 2019). In fact, *E. akaara* eDNA had an estimated half-life of 5.29 hours and reached concentrations below LOQ after two days of fish removal, indicating most fractions of eDNA can only persist for a limited time scale. Such short-term persistence further suggests eDNA as a sensitive method to provide contemporary snapshots that can detect the presence/ absence of target species confidently, which is coherent with previous studies on rapid eDNA degradation in marine environment (Collins et al., 2018; Murakami et al., 2019).

We acknowledge the fact that with only one system of experimental tank in this study, we cannot account for variability that may exist between different system set ups, such as microbial communities that could promote biotic degradation on eDNA (Hansen et al., 2018). Nonetheless, our work provides the first documented shedding and decay rate of a Serranid, *E. akaara* in a controlled experimental system, and offers valuable data to understand persistence of eDNA for application in monitoring.

Future directions should focus on disentangling the problems arise from geometric and metabolic variation on biomass estimation, where it was reported that juvenile or fish in developmental stage would shed more eDNA per biomass compare to adults due to higher total surface area to volume ratio as well as metabolic rates as previously seen in brook trouts and bluegill sunfish (Maruyama et al., 2014b; Yates et al., 2021), which could lead to erroneous estimation on fish biomass. It is also fundamental to understand how eDNA production and degradation of *E. akaara* would co-vary with ecological factors such as fish density, and environmental parameters including temperature, turbidity and pH.

### Application of assay in environmental seawater samples

In this study we conducted extensive eDNA sampling both spatially and temporally, covering locations and habitats with previous *E. akaara* distribution records, as well as local marine protected areas where neither recreational nor commercial fishing is permitted. We also deployed sampling methods that can increase captured diversity thus maximizing the likelihood of detecting this locally rare species, including sampling seawater along a 2km coastline transect, and large filter volume instead of several replicates of smaller volume (Ahn et al., 2022; Mathon et al., 2023). Despite the efforts, we could barely detect target species eDNA in the environmental seawater samples with qPCR, with only 6 samples (6.82%) in total showing minimal traces of target species eDNA. Although we did not conduct simultaneous fishery/ underwater visual census (UVC) survey to account for our qPCR results, we believe the results reflect the current *E. akaara* population status in Hong Kong. This is supported by a decade-long local reef fish UVC dataset with more than 7,000 man-hours underwater, where less than 10 individuals of *E. akaara* were observed between 2014 and 2023 (144°E Hong Kong Reef Fish Survey, 2023). We cannot rule out the possibility of false negative results on presence of low biomass i.e. one individual of *E. akaara*, as previous qPCR assays on black seabass, winter flounder and summer flounder could not detect the target species when only one individual was presented (as indicated by bottom trawling data) in the surveyed area of large water body (Kirtane et al., 2021). Nonetheless, our results still imply the population or biomass of *E. akaara* across Hong Kong and seasons are extremely low providing the same overall conclusion on the diminished population of *E. akaara* in Hong Kong.

We detected minimal traces of *E. akaara* eDNA at 4 sampling sites, which were Kung Chau (KC), Che Lei Pai (CLP), Cape d’aguilar (SWIMS) and Ninepin Island (NP). All four locations are documented with the presence of *E. akaara* in previous surveys or historical records (Cornish, 2000; Liu & Sadovy De Mitcheson, 2009; Sadovy & Cornish, 2000; 114°E Hong Kong Reef Fish Survey, 2023). We determined the risk of false positives from contamination for these samples would be minimal. Other than no amplified signals from negative controls, the sampling sites are fairly remote with considerable distances from local seafood restaurants or markets with live *E. akaara*, and therefore unlikely to have contamination from wastewater discharge. While there is a floating raft aquaculture farm within 2km radius of CLP, the farm exclusively cultures pearl oyster and the operation had been halted during time of sampling (Yan et al., 2019), thus pose little contamination threats to our sampling. On the other hand, there were only 1-2 amplifications out of 5 qPCR technical replicates in positive samples. The minimal amount of target sequence in the sample likely accounts for the low detection rate across qPCR technical replicates instead of PCR inhibition, as we deployed prefilters to remove large particle PCR inhibitors and extracted DNA of high quality and quantity with the modified protocol. Our results underscore the importance of including more technical replicates to reduce false negatives, especially when target sequences are at extremely low concentrations, such as for rare or endangered species, or samples with severely degraded DNA like ancient eDNA.

While all detected eDNA concentration of positive qPCR technical replicates were below LOD of the assay, detecting eDNA signals at low concentrations still holds major significance, as it still contains valuable information about the presence of target sequences even in trace amounts below desired confidence level of detection, and the data should not be disregarded (Klymus et al., 2020). As our data fulfilled all three criteria of detection below LOD to be considered as true positives (Klymus et al. 2020), which are detection made below 40 qPCR cycles, uniform amplification curve morphology and no amplification in negative controls, we regard these results as true positive qualitative data which indicates *E. akaara* presence at the sites.

### Implication for conservation and monitoring

With only few positive records of *E. akaara* across spatial and temporal sampling, we provide evidence that the population of *E. akaara* in Hong Kong is facing severe risk of local extirpation, showing no sign of recovery from the drastic population decline since 1970s due to overfishing (Liu & Sadovy De Mitcheson, 2009). The only current management on *E. akaara* i.e. marine protected areas only extends across 3.7% of total territorial waters of Hong Kong and is shown to be inadequate for conserving the remaining populations, as only 1 out of 4 sampling sites with positive eDNA records was a marine protected area (SWIMS; Cape d’Aguilar Marine Reserve).

Previous studies had detected *E. akaara* eDNA throughout its distribution range with eDNA metabarcoding, including East China Sea, Yellow Sea and the Sea of Japan (Ahn et al., 2020; Kenji et al., 2022; Kim, 2020; Lin et al., 2022). However, no eDNA metabarcoding study in the South China Sea ever recorded the presence of *E. akaara*, especially in Guangdong Province of China where *E. akaara* fishery was once prevalent (Cheang et al., 2020; Liu & Sadovy De Mitcheson, 2009; Zhang et al., 2024; Zhou et al., 2022; Chung et al., 2024). Hence, we suggest *E. akaara* in Hong Kong and South China Sea region as a whole could be currently suffering the worst population devastation compared to other populations across its distribution range.

We demonstrate that qualitative data from qPCR assays, indicating presence or absence of species can be a valuable resource for understanding the current distribution and habitat of threatened species, even when the target species are present in exceptionally low abundance. This information can be used to assess the effectiveness of existing conservation and management efforts. To improve the presence/absence based eDNA detection in future studies, site occupancy models can be incorporated into data analyses in the future to test how detection probability could vary in different target species densities, and further examine on the impacts of environmental covariates on occurrence and detection probabilities (Strickland & Roberts, 2019; Uthicke et al., 2022).

Overall, we have developed a reliable, species-specific qPCR assay for the detection of eDNA from the endangered species *E. akaara,* featuring with high sensitivity and specificity. Furthermore, we have generated new data on shedding and decay rates from tank experiments for these heavily fished and commercial targeted taxa within the Serranidae family. These findings have significant potential applications in biomass estimation, fishery stock management, and evaluation of current and future species enhancement initiatives for these species. The low detection rate of *E. akaara* across various spatial and temporal contexts highlights the serious risks of local extirpation of this species with no evident signs of recovery. This also demonstrates the capability and potential of using eDNA-based methodology to detect rare and endangered species in dynamic marine environment.

## DATA AVAILABILITY

The raw qPCR Cq values for assay validation and experiments are provided in supplementary materials.

## ACKNOWLEGEMENTS

We would like to thank boat skippers Ivan Kwok, Vincent Tong and Max Leung with their assistance in optimizing the on-site filtering set up. Thank you Miko Lui, Jacky Wong and other Ocean Park husbandry staffs on making the experiment possible. Thank you Yiu Wai Hong from 114°E Hong Kong Reef Fish Survey for providing underwater photograph to this manuscript. We also appreciate the feedback and support from the Schunter lab.

