## Supplementary Figures and tables for "Novel assay for endangered Hong Kong grouper (*Epinephelus akaara*) to assess eDNA shedding, decay, and population status"

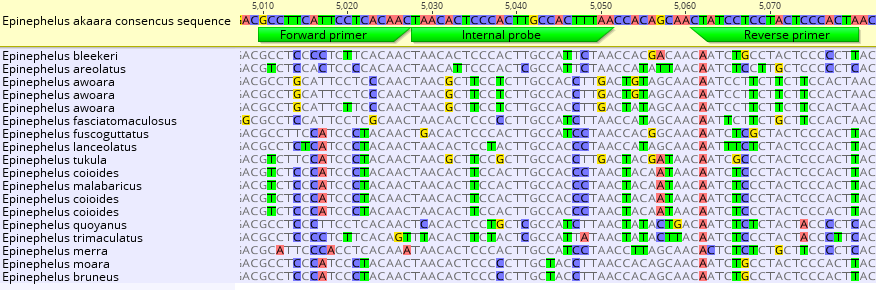

**Supplementary Figure 1.** Alignments and mismatches of base pairs between *E. akaara* reference consensus mitochondrial sequences and mitochondrial sequences from non-target congeneric species that can be found within the coastal environment of Northern South China Sea alongside *E. akaara*.

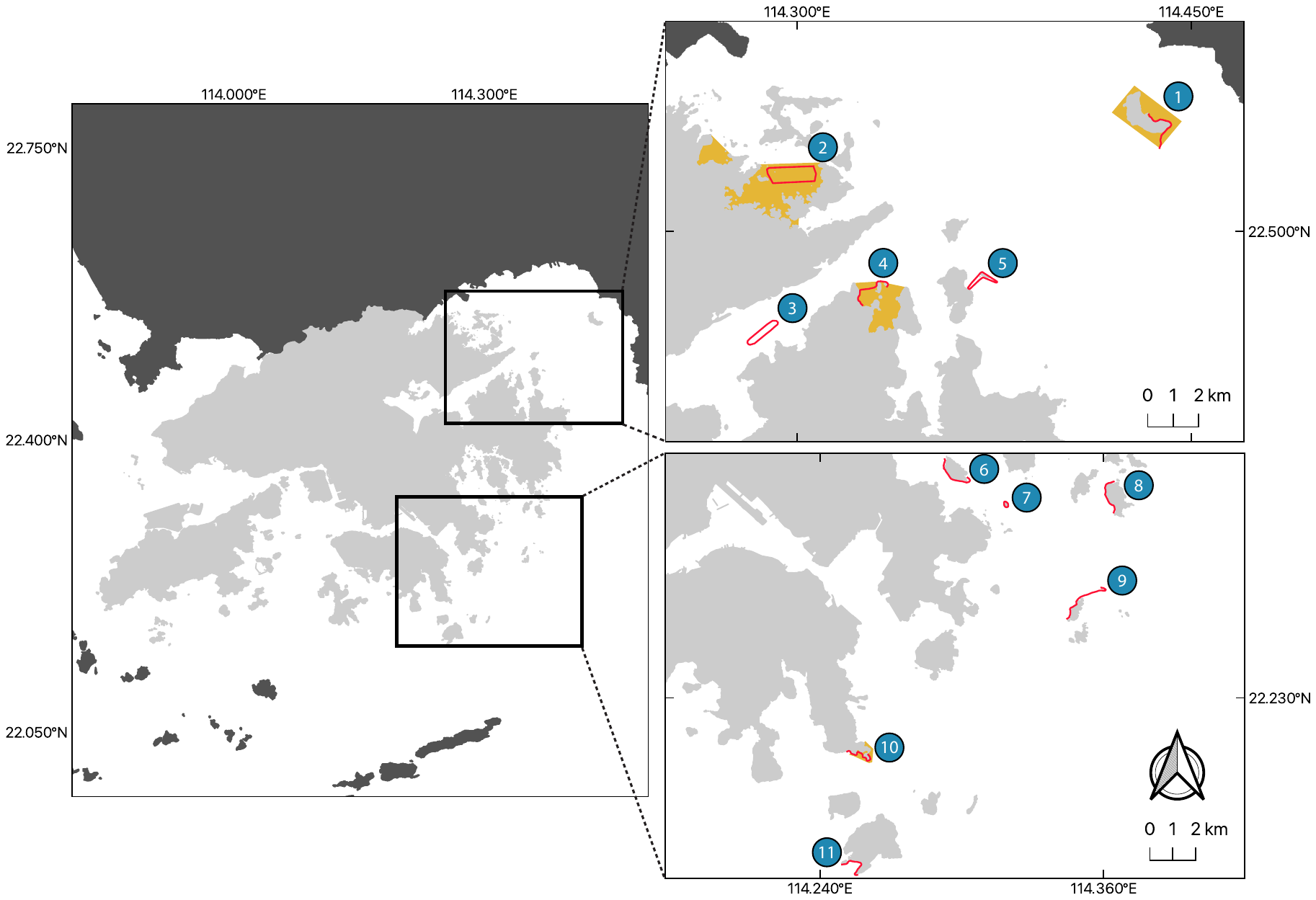

**Supplementary Figure 2.** Sampling locations across Eastern Hong Kong. Red lines indicated the water sampling transect for eDNA filtration at each site. Yellow area indicated current marine protected areas. Numbers in blue circles represented the 11 different sampling sites, namely: 1. Tung Ping Chau Marine Park; 2. Yan Chau Tong Marine Park; 3. Che Lei Pai; Hoi Ha Wan Marine Park; 5. Kung Chau; 6; Shelter Island; 7. Table Island; 8. Basalt Island; 9. Ninepin Island; 10. Cape d’Aguilar Marine Reserve; 11. Po Toi

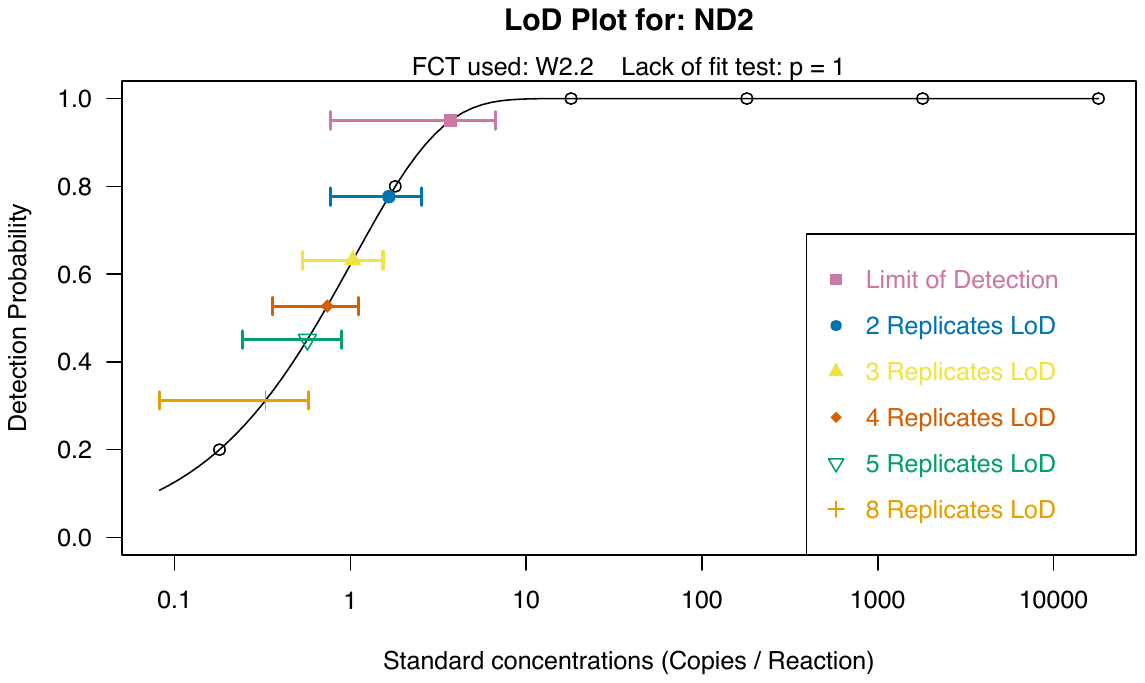

**Supplementary Figure 3.** LOD plot with detection probability as y axis, and standard concentrations as x axis. The solid line represents the LOD model determined by the calculator script from Klymus et al. (2020), while points of open circle indicate the detection rates of each standard tested. LOD is shown in the figure with pink square, and error bars represent 95% confidence intervals. Effective LODs were also drawn for multiple replicate analyses.

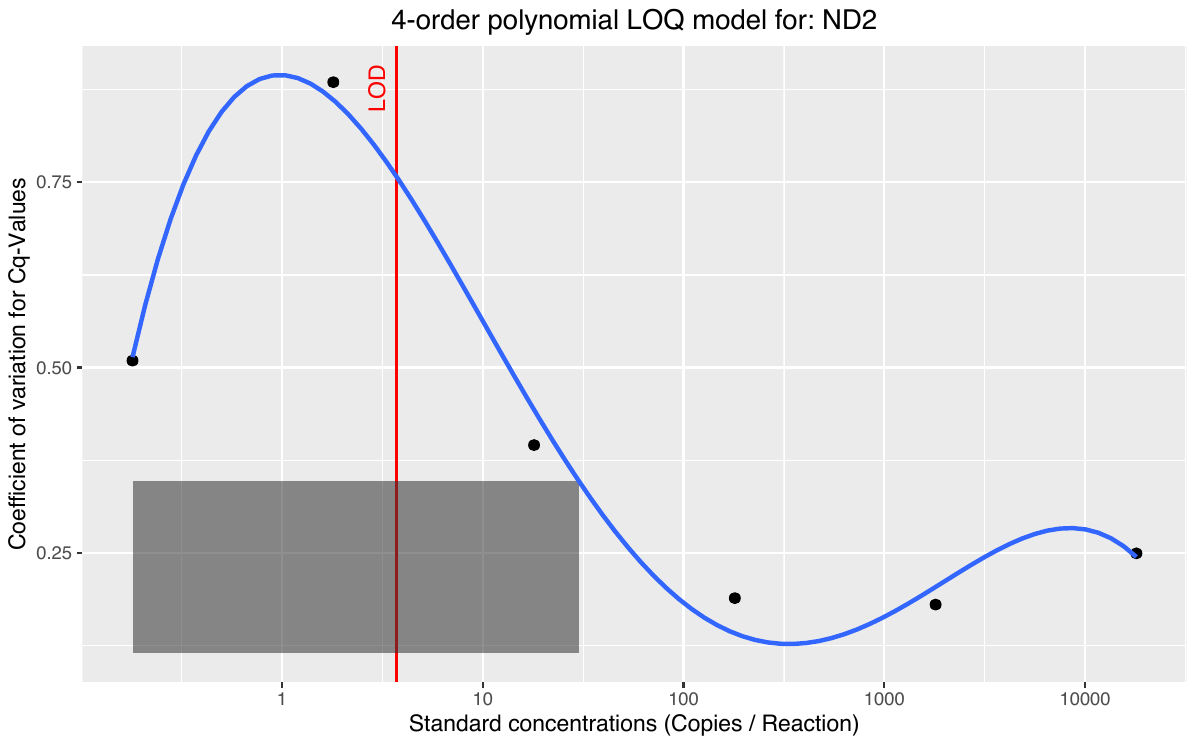

**Supplementary Figure 4.** LOQ plot with Coefficient of variation (CV) as y axis and standard concentration as x axis. Blue line represents the LOQ model determined by the calculator script from Klymus et al. (2020). The upper limit of grey rectangle was set as 0.35, which is the level of CV determine LOQ as defined in this study. The intersection between blue line and grey rectangle is the calculated LOQ. Vertical red line is the calculated LOD for reference.

Supplementary Table 1. Water parameters (Mean ± standard deviation) of tank across the experiment

| Experiment | Dissolved Oxygen (%) | pH | Temperature | Salinity | Water volume (L) |
| --- | --- | --- | --- | --- | --- |
| E.akaara | 95.1 ± 1.03 | 7.97 ± 0.07 | 26.31 ± 0.53 | 33.6 ±0.62 | 680 |

Supplementary Table 2. Non target close-related or co-occurring congeneric species that can be found within the coastal environment of Northern South China Sea alongside E.akaara.

**Species**

*Epinephelus aerolatus*

*Epinephelus awoara*

*Epinephelus bleekeri*

*Epinephelus bruneus*

*Epinephelus fasciatus*

*Epinephelus fasciatomaculosus*

*Epinephelus quoyanus*

*Epinephelus maculatus*

*Epinephelus merra*

*Epinephelus trimaculatus*
